# Single point mutations can potentially enhance infectivity of SARS-CoV-2 revealed by in silico affinity maturation and SPR assay

**DOI:** 10.1101/2020.12.24.424245

**Authors:** Ting Xue, Weikun Wu, Ning Guo, Chengyong Wu, Jian Huang, Lipeng Lai, Hong Liu, Yalun Li, Tianyuan Wang, Yuxi Wang

**Author notes:** These authors contributed equally to this work (Ting Xue, Weikun Wu). **Corresponding author**, Correspondence to Tianyuan Wang and Yuxi Wang. Email Address.

## Abstract

The RBD (receptor binding domain) of the SARS-CoV-2 virus S (spike) protein mediates the viral cell attachment and serves as a promising target for therapeutics development. Mutations on the S-RBD may alter its affinity to cell receptor and affect the potency of vaccines and antibodies. Here we used an *in-silico* approach to predict how mutations on RBD affect its binding affinity to hACE2 (human angiotensin-converting enzyme2). The effect of all single point mutations on the interface was predicted. SPR assay result shows that 6 out of 9 selected mutations can strengthen binding affinity. Our prediction has reasonable agreement with the previous deep mutational scan results and recently reported mutants. Our work demonstrated *in silico* method as a powerful tool to forecast more powerful virus mutants, which will significantly benefit for the development of broadly neutralizing vaccine and antibody.

## Introduction

Since it first broke out in late 2019, COVID-19 has soon spread out worldwide and has been defined as a world pandemic by WTO. The disease is caused by a novel coronavirus SARS-CoV-2 (Severe Acute Respiratory Syndrome Coronavirus 2), which is a beta-coronavirus closely related to the known SARS-CoV^1 2 3^. Much efforts have been distributed in developing the prevention/treatment to the disease, including small molecule drug^4^, vaccine^5^, neutralizing antibodies^6–7^ and other engineered proteins ^8–9^. However, none of the methods has been comprehensively tested or publicly applied so far.

The adhesion of the virus to the target cell and the following membrane fusion process are essential steps in virus infection thus those two processes are promising targets for drug development. As is typical for coronavirus, the homo-trimeric spike glycoprotein (S protein, comprising S1 and S2 subunit in each monomer) on the envelope of SARS-CoV-2 is responsible for the cell adhesion process. SARS-CoV-2 uses hACE2 as the receptor for host cell entry^10–11^, and the dissociation constant *K*_D_ of S protein RBD binding to hACE2 was determined to be in the scale of nM^10–12^. X-ray crystal structures of RBD had been solved in complex with the hACE2 receptor, revealing essential interactions on the binding surface^13–14^.

As cell entry is the very first step of virus infection, blocking the binding of S protein to hACE2 can potentially prevent virus transmission. Many developing therapeutics are based on this concept. Monoclonal neutralizing antibodies separated from convalescent patient showed complete competition with RBD and can reduce virus titers in infected lungs in a mouse model^7^. Recombinant vaccine that comprises residues from the S protein RBD was shown to induce functional antibody response in immunized animals^15^. What’s more, de novo protein inhibitors with *K*_D_ of picomolar-level has been designed to neutralize the virus by targeting S protein RBD^8–9^.

While therapeutics relying on the RBD binding surface require the surface to be consistent enough, SARS-CoV-2 is an RNA virus that is known to have high mutation rate. According to reported data, 196 missense mutations in the gene encoding the RBD domain have been discovered^16^. Although most mutants on the RBD domains were determined to be less infectious, some variants indeed became resistant to neutralizing antibodies^17^. We are interested in finding RBD mutants with higher affinity, which may escape future treatments.

*In-silico* affinity maturation technology is widely used in protein engineering and antibody discovery^18^. Rosetta Flex ddG method is a ddG estimation method developed within the Rosetta macromolecule modeling suite. The ddG represents the difference in protein-protein interaction strength upon mutation. The protocol generates an ensemble of models to include conformational plasticity around a specific/given mutation site and then calculates the average ddG over the ensemble. This method has been shown to outperform previous methods, and considerable improvement on predicting binding-stabilizing mutations was observed^19^.

In this article, Rosetta Flex ddG protocol was used to predict the binding affinity change of point mutations on the RBD binding surface. Candidate mutants with large negative predicted ddG score were selected for further experimental validation. 6 of the 9 recommended mutants shows improved affinity to hACE2 in SPR affinity assay.

## Results and Discussion

### Mutations with increased binding affinity are accurately predicted through Rosetta Flex ddG and validated by SPR assay

Rosetta Flex ddG calculation was carried out for 25 residues on the RBD binding surface. For each residue, the wild type (WT) amino acid was mutated to the other 19 natural amino acids individually and Flex ddG calculation was applied. 48 models for each mutant were generated and the ddG score were averaged over 48 models. In all 475 possible single point mutants (25 times 19), 114 mutants have negative ddG score, which implies potential mutations with improved affinity. The lowest predicted ddG score was −3.66 REU (Rosetta Energy Unit) from mutation Q498W. Mutants with large negative ddG values were examined manually and 9 mutants were selected for SPR (Surface Plasmon Resonance) assay. The selected mutants and their corresponding predicted ddG scores are shown in **Table1**.

**Table 1:** Results of the binding affinity predicted by Rosetta Flex ddG and measured by SPR assay.

| Mutant | Flex ddG-$\Delta\Delta G_{\text{binding}}$<sup>a</sup> | SPR- $K_D$<sup>b</sup> | SPR-$\Delta\Delta G_{\text{binding}}$<sup>c</sup> |
| --- | --- | --- | --- |
| WT | 0.00 | 21.08±3.01 | 0.00 |
| Q498W | -3.66±1.80 | 7.10 | -2.70 |
| Q498R | -2.04±1.34 | 11.60 | -1.48 |
| T500W | -1.90±0.56 | 21.80 | 0.08 |
| S477H | -1.39±1.16 | 13.90 | -1.03 |
| Y505W | -1.23±0.41 | 16.10 | -0.67 |
| T500R | -1.21±1.38 | 12.20 | -1.36 |
| N501V | -1.02±1.09 | 158.50 | 5.00 |
| Y489W | -1.01±0.50 | 38.90 | 1.52 |
| Q493M | -0.82±1.39 | 6.90 | -2.77 |

<sup>a</sup> $\Delta\Delta G_{\text{binding}}$ calculated by Rosetta Flex ddG. dG\_cross is used as ddG score, which means the binding energy of the interface calculated with cross-interface energy terms. The unit is REU. Uncertainties were estimated from the standard deviations of 48 replicated trajectories.
<sup>b</sup> $K_D$ (dissociation constant) assayed by SPR. The unit is nM. For WT, uncertainties were estimated from the standard deviations of 3 replicated experiments. For other mutants, the experiments were carried out only once.
<sup>c</sup> $\Delta\Delta G_{\text{binding}}$ calculated from $K_D$ measured by SPR. The unit is kcal/mol.

Next, we carried out SPR affinity measurement for WT RBD and 9 potentially stabilizing mutants to validate the prediction result. **Figure 1** shows the SPR sensorgram of WT and the mutant Q493M. For other mutants, the SPR sensorgram are shown in **Figure S4**. The experimental *K*_D_ are reported in **Table1**. 6 of the 9 selected mutants with predicted elevated binding affinity have lower *K*_D_ to hACE2 than WT RBD in SPR assay. The prediction accuracy is comparable to that reported in previous work^19^. Among the 9 selected mutants, the best binder is mutation Q493M with a *K*_D_ of 6.90 nM, showing a 3-fold improvement in affinity compared with that of WT. There is a modest Pearson correlation coefficient of 0.41 between predicted ddG and SPR measured *K*_D_ (**Figure S2**).

**Figure 1.**
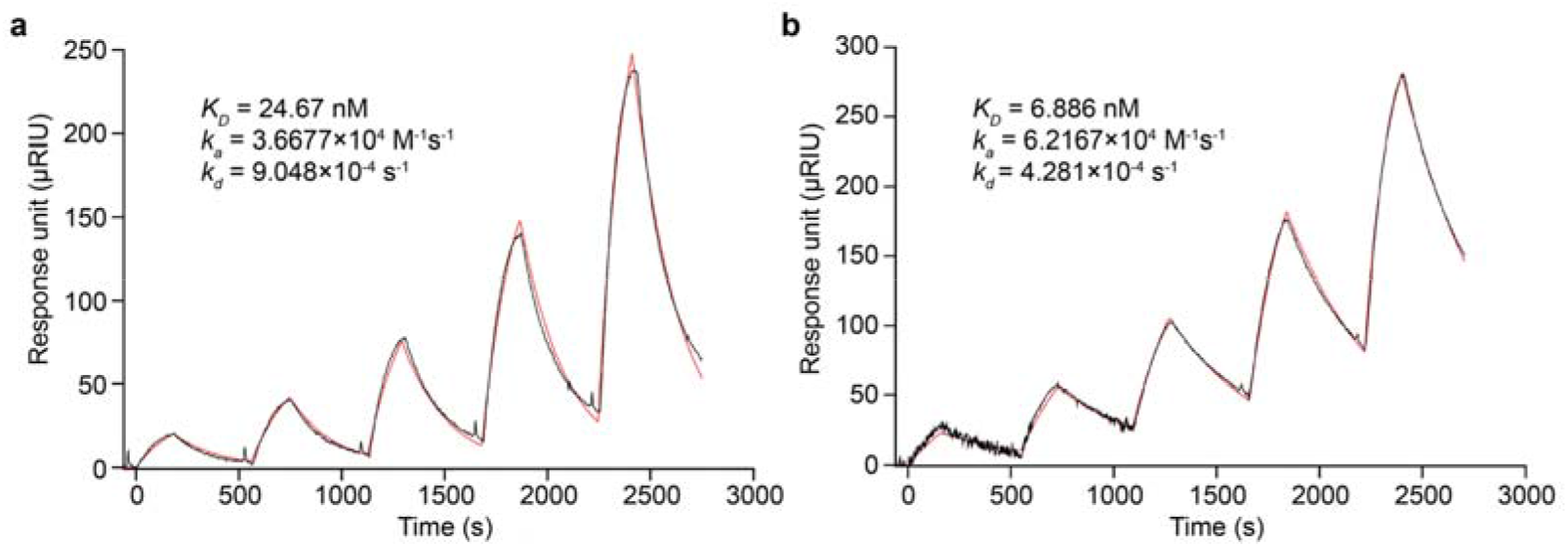
Surface plasmon resonance sensorgram showing the binding kinetics for human hACE2 and immobilized SARS-CoV-2 S protein RBD (a) wildtype / (b) mutant Q493M. Data are shown as black lines, and the best fit of the data to a 1:1 binding model is shown in red lines.

Among the 5 RBD sites (477, 493, 498, 500 and 505) which give affinity-increased mutants, 4 of them were identified as key residues interacting with hACE2 (493, 498, 500 and 505)^14^ and 3 of them were SARS-CoV-2 unique mutants different from SARS-CoV-1 (477, 493 and 498)^13^ in previous research. The increased affinity coupled with those mutations shows that the key residues in natural binding pattern aren’t necessarily optimized for binding. And even after evolution towards more potent binding, there is still great capacity for increased binding affinity.

### Analysis of mutant structures indicates the stabilizing interaction mainly come from newly formed hydrogen bonds and hydrophobic interactions

To explain the affinity-increase effect in mutants, we analyzed the interface characters other than dG_cross reported by Rosetta InterfaceAnalyzeMover. We chose to analyze the hbond_int (count of hydrogen bonds crossing the interface) and the dSASA (buried solvent accessible area at the interface, Å^2^) as those terms are closely related to dG_cross value. In **Table 2**, each term was calculated by averaging over the 48 difference between the reported values of the WT model and the mutant model.

**Table2.** Interface feature differences calculated from Rosetta InterfaceAnalyzer

| mutant | mut_ddG | SPR result | hbonds_int | dSASA_int | dSASA_hphobic | dSASA_polar |
| --- | --- | --- | --- | --- | --- | --- |
| Q498W | -3.66 | stabilize | 0.31 | 39.61 | 58.39 | -18.78 |
| Q498R | -2.04 | stabilize | 1.65 | 27.11 | -8.08 | 35.20 |
| S477H | -1.39 | stabilize | 0.77 | 52.13 | 17.81 | 34.32 |
| Y505W | -1.23 | stabilize | 0.02 | 16.04 | 31.05 | -15.01 |
| T500R | -1.21 | stabilize | 0.71 | 43.28 | -5.05 | 48.33 |
| Q493M | -0.82 | stabilize | -0.92 | -4.67 | 71.23 | -75.90 |
| T500W | -1.90 | destabilize | -0.08 | 48.38 | 34.08 | 14.30 |
| N501V | -1.02 | destabilize | 0.00 | 0.05 | 12.30 | -12.25 |
| Y489W | -1.01 | destabilize | 0.04 | 49.17 | 50.73 | -1.56 |

hbonds\_int: the change of hydrogen bond numbers at the interface.
dSASA\_int: the change of solvent accessible area buried at the interface, in square Angstroms.
dSASA\_hphobic: the change of the hydrophobic part of solvent accessible area buried at the interface, in square Angstroms.
dSASA\_polar: the change of the polar part of solvent accessible area buried at the interface, in square Angstroms.

In the crystal structure, the WT Q498 forms a new hydrogen bond with nearby hACE2 42N (**Figure 2a**). According to the best Q498W mutant model, the 498W residue forms a hydrogen bond with hACE2 38D (**Figure 2b**). The Q498W mutation also increases hydrophobic contact with nearby residues, as indicated by the increment of hydrophobic SASA surface. For the Q498R mutant, in most of the mutant models, 498R forms hydrogen bond with hACE2 42N (**Figure 2c**), and in some of the models also with hACE2 38D (**Figure 2d**). This phenomenon could be explained by the fact that the longer hydrophobic part of arginine compared with glutamine could afford increased hydrophobic interaction. On average, there are 1.65 additional interface hydrogen bonds formed after mutation which may contribute to the increasing affinity. S477H mutation forms a new hydrogen bond with the terminal NH_2_ group of hACE2 19S (**Figure 2e**) according to the mutant model. Calculation result shows that additional 0.77 hydrogen bond formed in average along with increased hydrophobic interaction. For the Y505W mutant, the model shows hydrogen bond has no contribution. In the original crystal structure of WT, 505Y doesn’t form any hydrogen bond with other residues. The mutation Y505W is likely increasing the affinity by evolving to bear better hydrophobic contact (**Figure 2f**), as shown in the increase of hydrophobic interactions. For the T500R mutant, the dominant factor is newly formed hydrogen bond interaction which is similar to Q498R mutant. The WT 500T forms a hydrogen bond with 41Y (**Figure 2g**). In the best mutant model, T500R forms hydrogen bond with hACE2 329E and 330N (**Figure 2h**), while in some other model T500R only forms hydrogen bonds to 330N (**Figure 2i**). The Q493M mutant is the only affinity-increase mutant with significant decrease in hydrogen bond count. 493Q in WT originally forms a hydrogen bond with hACE2 35E (**Figure 2j**). The van de vaals contact increases significantly when mutating to a hydrophobic residue like methionine (**Figure 2k**), indicated by the large dSASA_hphobic in **Table 2**.

**Figure 2.**
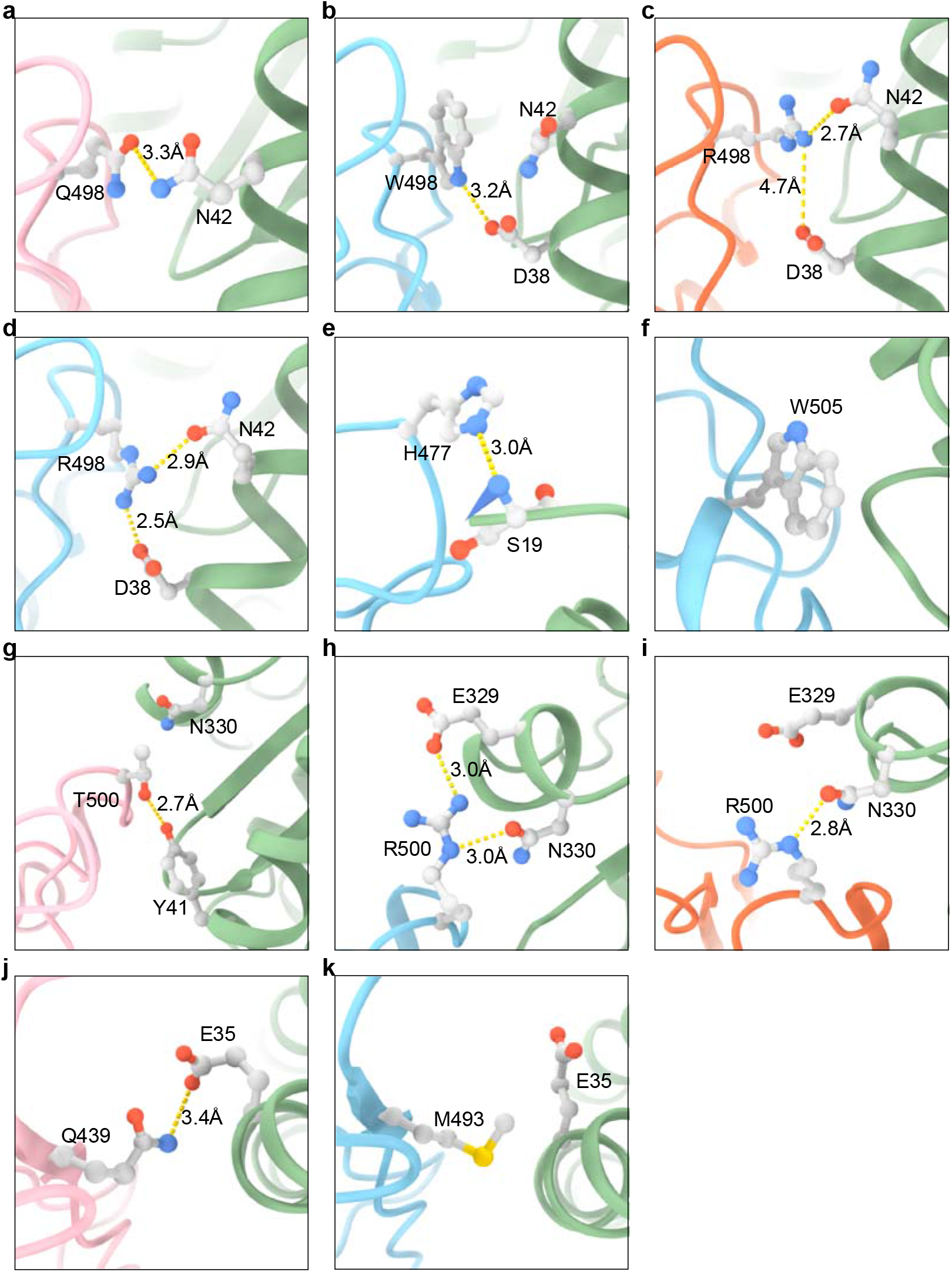
Predicted structure of mutants with enhanced affinity. The RBD of S protein and hACE2 are shown in cartoon representation. Residues of interest are shown in stick. Hydrogen bond is shown in yellow dash line. The distance between hydrogen bond donor and acceptor atoms is shown.

According to above analysis, the enhanced interaction mainly come from additional hydrogen bonds and hydrophobic interactions. The Q493M mutant also reveal that the hydrophobic term itself can be sufficient for leaning the equilibrium to more potent binding.

### The false positive predictions are possibly due to inaccurate estimation of hydrogen bond interaction

3 of the 9 mutants (T500W, N501V and Y489W) are predicted to stabilize the interaction, however according to the SPR experiment result they are neutral or destabilizing. Next, we try to analyze the Flex ddG energy terms and structure model to elucidate the reason of the incorrect prediction. For the mutant T500W, the increase of SASA_hphobic is the dominant factor to binding. However, the mutation to W destroy the hydrogen bond between 500T and hACE2 41Y as shown in T500R (**Figure 2g, Figure 3a**). Although in most of the WT model this hydrogen bonding pattern was well preserved, the hydrogen bond count didn’t show this adverse factor as significant, with only a decrease of 0.08. This probably leads to wrong estimation of binding affinity. For the mutant N501V, in the crystal structure of N501 WT, 501N adopted the rotamer with the NH_2_ of the amide pointing at hACE2 41Y, which forms a hydrogen bond (**Figure 3b**). However, in most WT structure model generated after backrub (a method to perturbate protein backbone to generate an ensemble of conformations), the oxygen atom of the amide points to 41Y instead with an O-O distance of 3.6 Å (**Figure 3c**). Therefore, the ddG calculation did not consider the destruction of the hydrogen bond, which is verified in the zero hbond count difference. For the mutant Y489W (**Figure 3e**), the situation is similar to N501V. In the crystal structure, the hydroxy oxygen of 489W is within 3.5 Å of hACE2 83Y (**Figure 3f**). However, in most WT models generated by backrub, the distance is larger than 3.7 Å (**Figure 3g**), resulting in loss of consideration for this hydrogen bond impact in the final ddG score.

**Figure 3.**
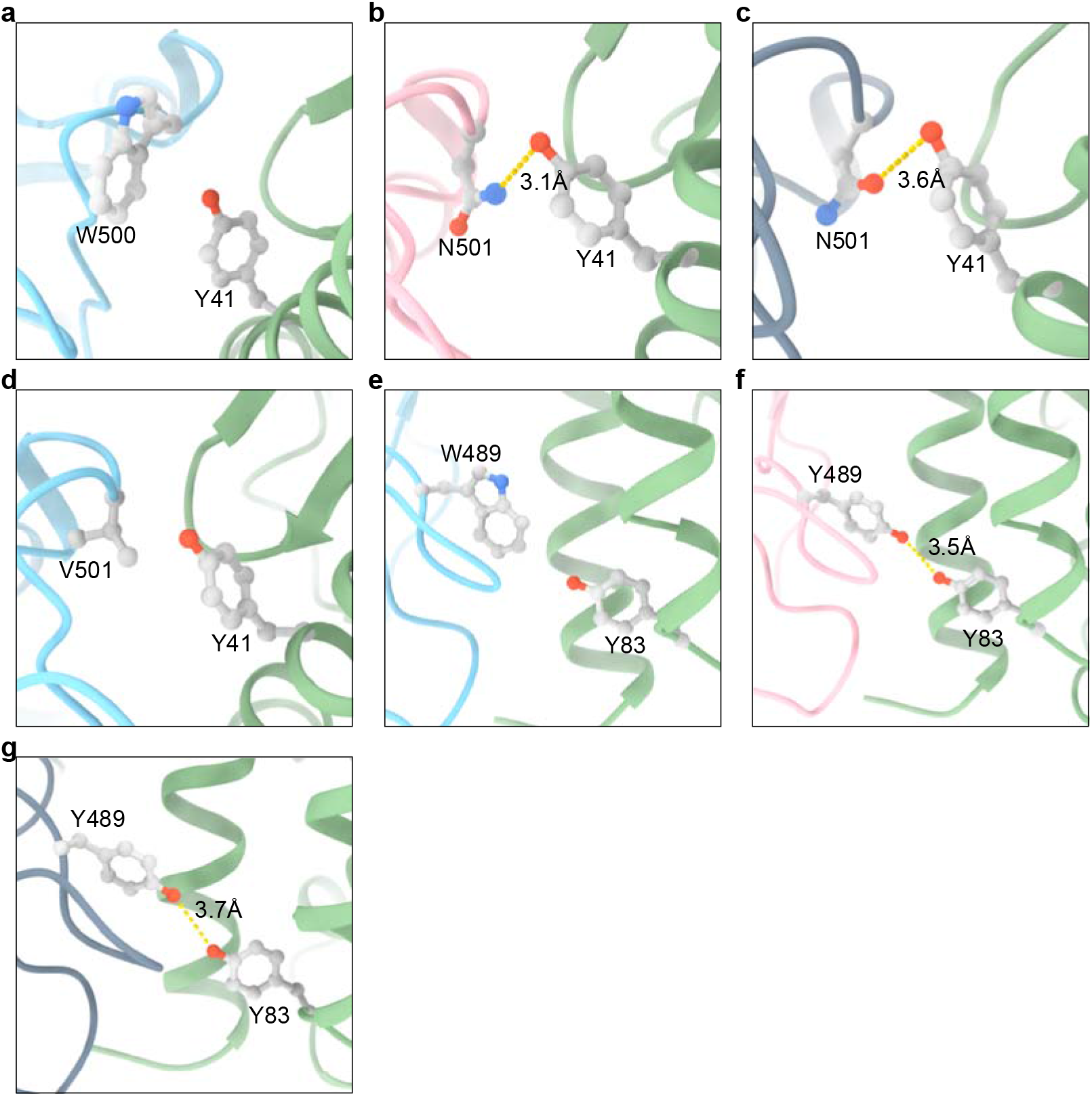
Predicted structure of mutants with reduced affinity. The RBD of S protein and hACE2 are shown in cartoon representation. Residues of interest are shown in stick. Hydrogen bond is shown in yellow dash line. The distance between hydrogen bond donor and acceptor atoms is shown.

By analyzing the models of false positive predictions, we illustrated that the main error may come from the broken of hydrogen bonds which are not considered in the ddG calculation. The missing hydrogen bond may be existing hydrogen bond in the scoring step and the hydrogen bond destroyed in the previous relaxing step. Visual examination of the model structure to see whether the original hydrogen bond is disrupted and comparing with the predicted surface properties may help spot those inaccuracies and exclude some false positive instance before wet-lab experiments.

### The predicted binding affinity modestly agrees with deep mutational scanning (DMS) results

Starr et al. has systematically changed every amino acid in the S protein RBD and determined the mutation effects on hACE2 binding using deep mutational scanning method^20^. Although this high throughput method using flow cytometry has some contradict results compared with our SPR results (**Figure S3**), it is nevertheless a valuable data set to evaluate our Flex ddG prediction. Plotting log (*K*_D,mut_/*K*_D,wt_) measured by DMS versus predicted ddG gives Pearson correlation coefficient of 0.37 (**Figure 4**).

**Figure 4.**
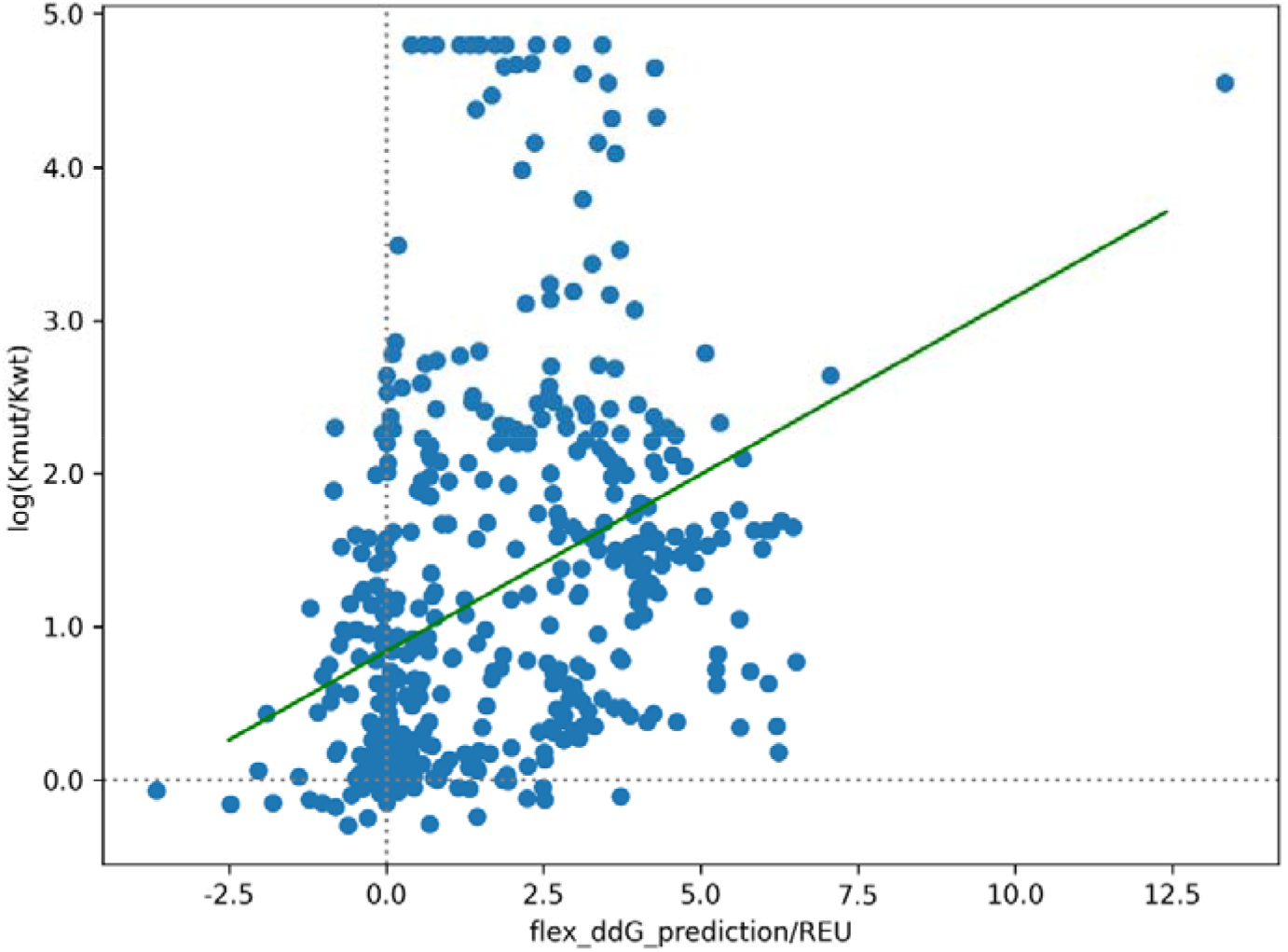
The log (*K*_d,mut_/*K*_d,mut_) from the DMS result versus our Flex ddG predicted ddG value.

Using the result measured by DMS as the ground truth, the prediction performance of Rosetta Flex ddG method is calculated. Remarkably, the precision on predicting the stabilizing mutation is 0.18, and the recall is 0.50 given a ratio of 40/432 stabilized and destabilizing mutations, which shows the potential of the Flex ddG method to suggest stabilized mutations for enhanced binding affinity (**Table S1**).

## Conclusion

In this article, we utilized the Rosetta Flex ddG protocol to predict mutations on the SARS-CoV-2 RBD binding surface that can strengthen its interaction to hACE2. SPR experiments confirmed 6 of the 9 predicted mutations had increased affinity, with the best mutation showed a 3-fold improvement of *K*_D_.

The SARS-CoV-2 mutants carrying the mutation Y453F were discovered in Denmark to circulate between human and minks and have brought up concerns recently^21–22^. Our prediction (−0.30 REU, supplementary material) and the deep mutational scan results^20^ both shows that the Y453F mutation will strengthen the interaction between RBD to hACE2, which is consistent with the maintained transmissibility and pathogenicity. Although more studies on the potential effect on treatment, diagnostic test and virus antigenicity are still ongoing, preliminary results showed that the Cluster-5 strain which carries another 3 mutations outside the RBD domains was more difficult to be recognized by convalescent sera^23^. It is also a concern that spreading of the virus in minks may bring up more fatal variants.

The still-mutating SARS-CoV-2 alerts us of the need of an efficient mechanism in dealing with newly discovered pathogens. For such RNA virus under constant mutagenesis, it is necessary to quickly spot possible mutants with increased pathogenicity or infectivity. Given the relatively low cost and short period of computational methods, we can build a computational forecast system on virus pathogenicity in the early stage of a virus breakout. The Flex ddG method used in this work would be a constructive part in this system, predicting pathogenicity by evaluating mutant binding affinity to cell receptor. The mutations on the binding surface are especially essential in vaccine or antibody development and the predicted mutations can be considered in designing multi valent vaccine to prevent possible immune escape.

Furthermore, *in-silico* affinity maturation itself is a promising technology in the development of macromolecule drugs. For instance, it can help reduce the scale of experimental screening in optimizing the affinity of peptide or protein drugs. Our work demonstrated the ability of the previously reported Flex ddG method to serve as an efficient affinity maturation method in predicting stabilizing mutants with no need for the time-consuming MD process. We anticipate that combined with machine learning and deep learning technology, Rosetta Flex ddG-based *in-silico* affinity maturation can be improved to be faster and more accurate in the near future.

## Methods

### Rosetta Flex ddG calculation

The S protein RBD-hACE2 complex structure was downloaded from PDB database (PDB ID 6M0J)^13^. The structure was relaxed using Rosetta FastRelax Mover. Resfiles describing saturate point mutation were generated for each residue on the S protein within 8 Å of S protein and hACE2 interface in the relaxed structure. The Flex ddG protocol defined in previous literature^19^ was refactored for in-house high-performance computing platform, and implemented using pyRosetta API (**Figure S1**). For each mutant defined by a resfile, backrub sampling was applied around the mutation site. The structure was then allowed to repack and relax globally with both the WT and the mutant. The binding energy dG_cross was calculated using the InterfaceAnalyzerMover, and the dG_cross difference between the mutant and the WT model was taken as ddG. 48 independent ddG calculations were perform for each mutant. The mutants were sorted according to their average ddG score. After subsequent manual examination of top scored structures, 9 structures were selected for further SPR wet-lab experimental validation.

### SPR assay

The affinity between SARS-CoV-2 Spike Protein (RBD, His Tag) and hACE2 was measured using a Reichert4SPR system (Reichert Technologies, Depew, NY, USA) in single-cycle mode. SARS-CoV-2 Spike Protein (RBD, His Tag) and its mutants were immobilized to an mSAM sensor chip (Planar Polyethylene Glycol/Carboxyl Sensor Chip P/N 13206061) at approximately 500 response units. The experiment data were obtained at 25°C with running buffer PBST (8 mM Na_2_HPO_4_, 136 mM NaCl, 2 mM KH_2_PO_4_, 2.6 mM KCl, and 0.05% (v/v) Tween 20, pH7.4). Gradient concentrations of hACE2 (from 100 nM to 6.25 nM with 2-fold dilution) were then flowed over the chip surface. The resulting data were fit to a 1:1 binding model using Scrubber and Clamp software. Kinetic rate constant *k_a_* and *k_d_* were calculated from the above analysis and the apparent *K*_D_ was calculated as *k*_d_ / *k*_a_.

### Plasmid construction, protein expression and purification

The receptor binding domain (RBD, corresponding to Spike 319-514AA) of spike protein of SARS-COV-2 have been previously codon optimized and synthesized into pTT5 vector (EcoRI + BamHI) through PCR based overlapping oligonucleotides assembly. Kozak sequence was inserted before start codon to enhance translational efficiency. To obtain the secreted RBD protein, a mouse V-set immunoregulatory receptor signal peptide was fused to the N-terminus of the protein (1-32, MGVPAVPEASSPRWGTLLLAIFLAASRGLVAA). Ten mutations were introduced using primers with the desired mutation in a PCR protocol which amplifies the entire plasmid template (the above clone as the template). Restriction enzyme digestion and Sanger sequencing confirmed that all 10 clones were correct.

The mutated RBD proteins were prepared using the Expi293 Expression System (Code: A14635). The specific operation was performed following the kit instruction. Expi293F cells were maintained in serum-free Expi293 Expression Medium, and the Expression plasmid transfected into Expi293 cells by using an ExpiFectamine 293 Transfection Kit (all from Gibco). Five days after transfection the medium was collected, and the protein was purified by Ni-NTA (QIAGEN) column chromatography (nonspecifically binding contaminants were washed on Ni-NTA column using PBS, pH 7.4 containing 20 mM imidazole and eluted with PBS, pH 7.4 supplemented with 300 mM imidazole.). The eluted fractions were pooled and dialyzed against PBS (pH7.4) to remove the imidazole. Purified proteins were checked by SDS-PAGE and protein concentrations were quantified using Nano-Drop.

## Data availability

Relevant data are within the paper and its Supporting Information files. Other data and parameters generated or analyzed during the study are available from the corresponding author on reasonable request.

## Funding

Part of this work was supported by Alibaba Cloud Anti-COVID-19 Program. Part of this work was supported by Special Funds for COVID-19 Prevention and Control of West China Hospital of Sichuan University (HX-2019-nCoV-003), Sichuan Science and Technology Program (2019YFS0003), Technological Special Project for ‘Significant New Drugs Development’ (2018ZX09201018-021), China Postdoctoral Science Foundation (2018T110984, 2017M610607).

## Author contributions

Tianyuan Wang and Yuxi Wang conceived the study and edited the paper. Ting Xue, Weikun Wu, Ning Guo, Chengyong Wu and Jian Huang conducted the experiments, performed the calculation and analyzed the results. Tianyuan Wang wrote the manuscript. All authors reviewed the manuscript.

## Acknowledgements

We gratefully thank Jingjia Liu for assisting in preparation of the manuscript.

## Competing interests

Weikun Wu, Ning Guo, Jian Huang, Lipeng Lai and Tianyuan Wang are employees of XtalPi - AI Research Center (XARC).

