## Supplementary file S1 for "Single point mutations can potentially enhance infectivity of SARS-CoV-2 revealed by in silico affinity maturation and SPR assay"

**Contents**

**Table S1.** The classification performance of flex ddG prediction result using DMS dataset as ground truth.

**Figure S1.** Schematic of the flex ddG protocol, modified from the flex ddG paper.

**Figure S2.** ddG calculated from SPR result versus flex ddG predicted ddG value.

**Figure S3.** The ddG calculated from SPR versus log(Kd,mut/Kd,wt) from the DMS paper.

**Table S2.** List of the sample name on the SDS-PAGE figure corresponding to the mutants.

**Figure S4.** Surface plasmon resonance sensorgram showing the binding kinetics for human ACE2 and immobilized SARS-CoV-2 S protein RBD wildtype / mutants.

**Figure S5.** SDS-PAGE result of the 10 spike protein RBD mutants.

**Table S3.** SPR result of RBD-WT and 10 mutants.

**Table S1. The classification performance of flex ddG prediction result using DMS dataset as ground truth.**

|  | DMS stabilizing | DMS neutral | DMS destabilizing | Precision |
| --- | --- | --- | --- | --- |
| Predicted stabilizing | 20 | 1 | 93 | 0.18 |
| Predicted neutral | 0 | 0 | 0 | 0 |
| Predicted destabilizing | 20 | 2 | 339 | 0.94 |
| Recall | 0.50 | 0 | 0.78 |  |

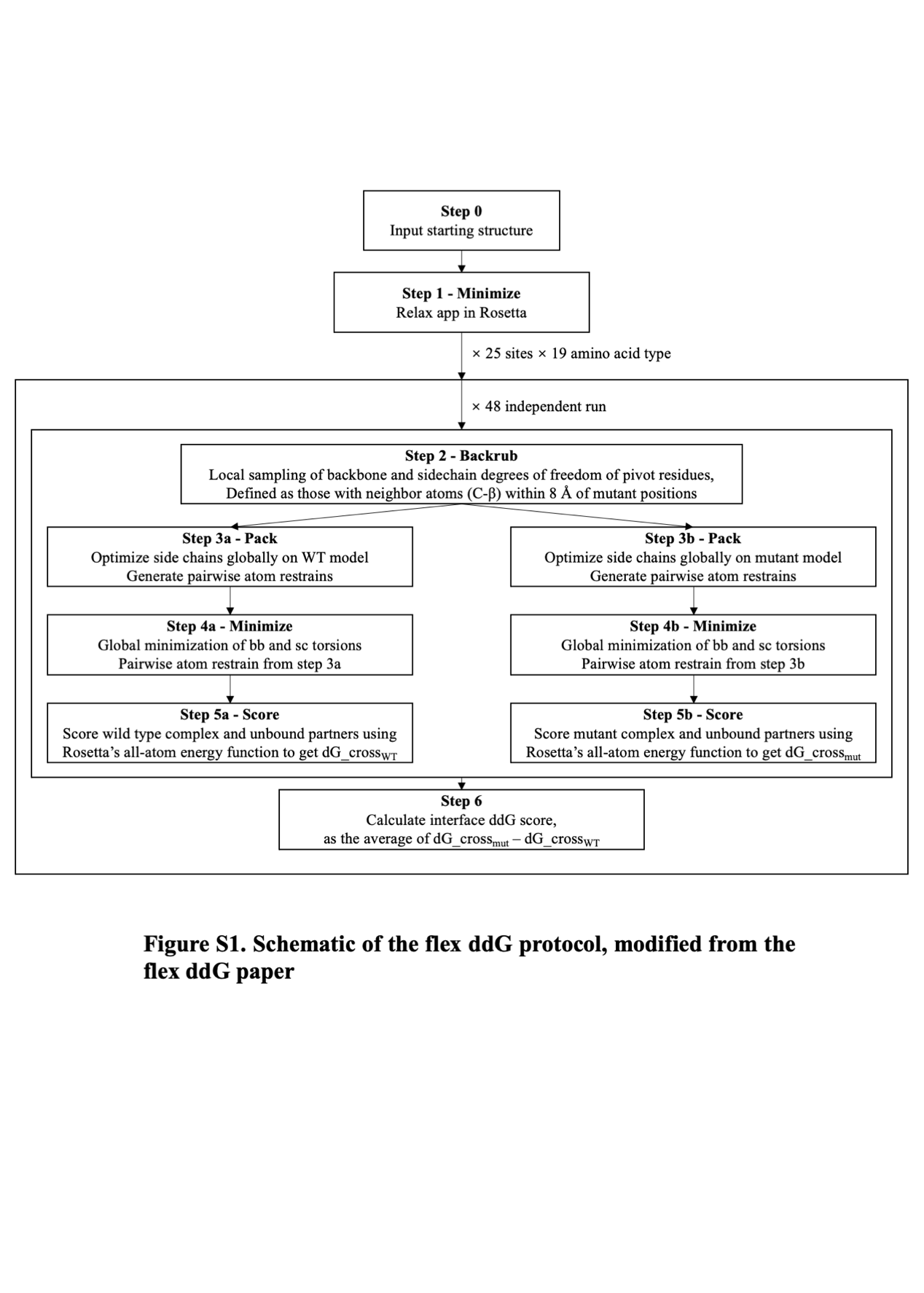

**Figure S1. Schematic of the flex ddG protocol, modified from the flex ddG paper.**

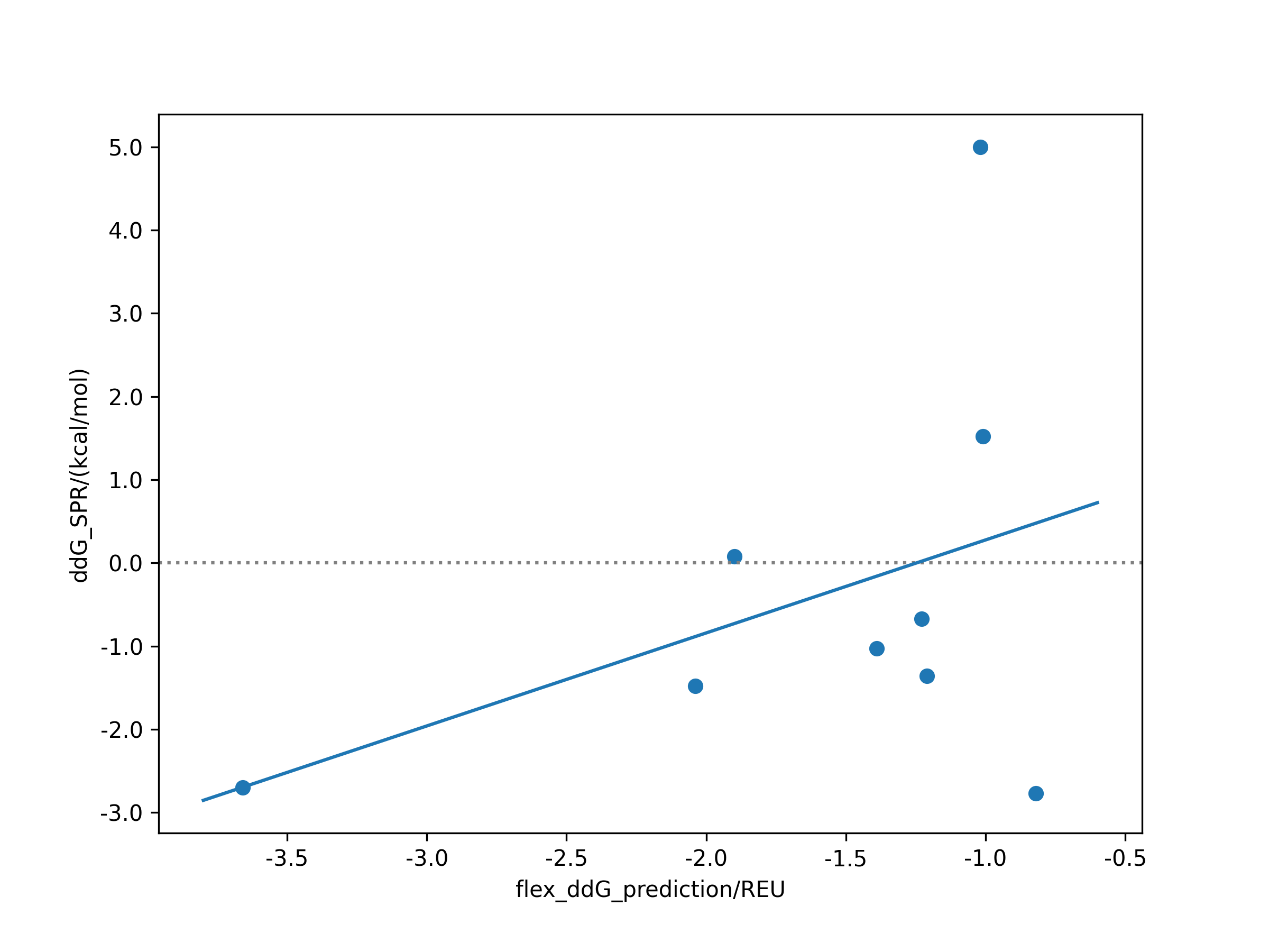

**Figure S2. ddG calculated from SPR result versus flex ddG predicted ddG value. The Pearson correlation coefficient of the regression is 0.41.**

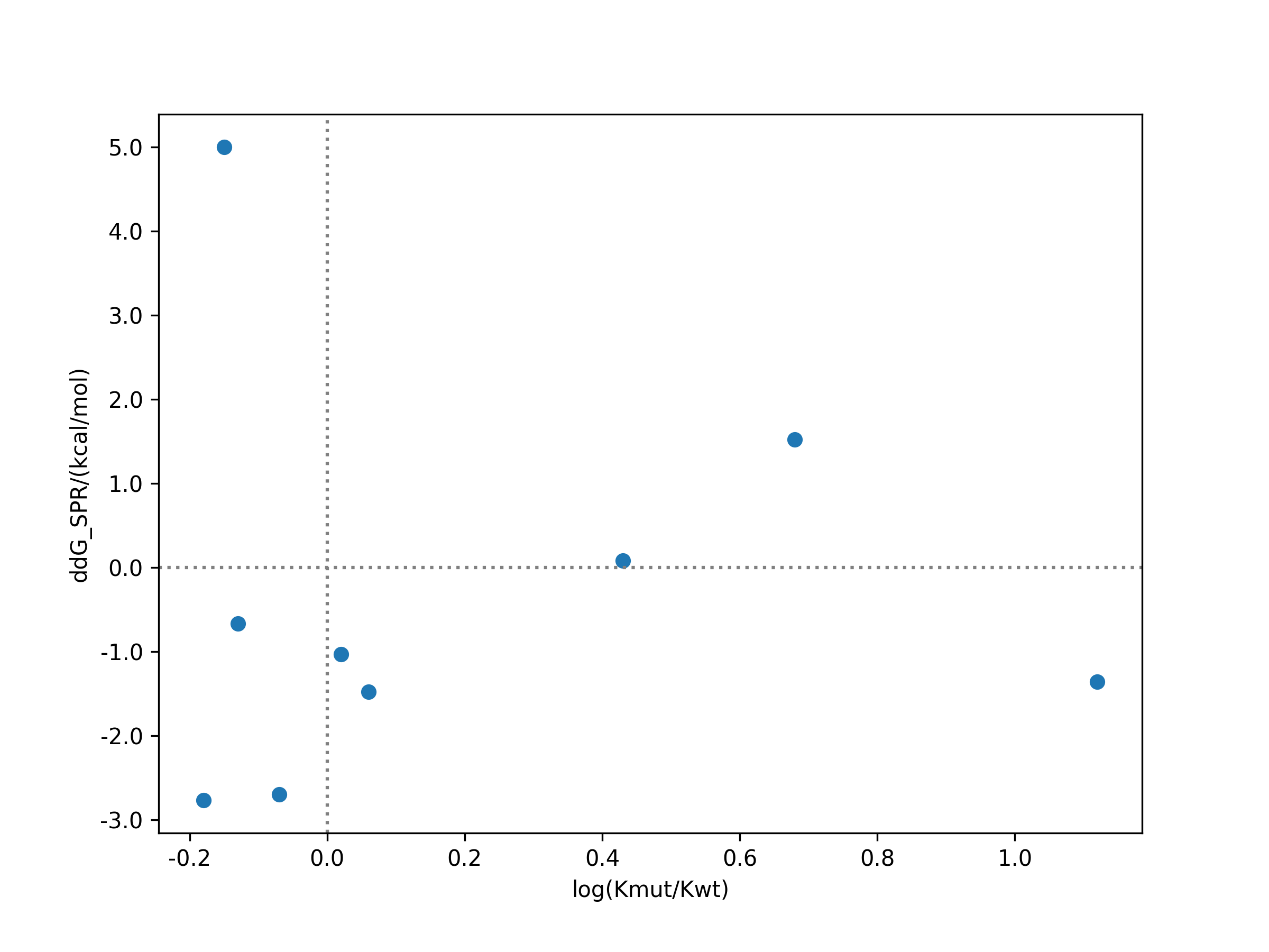

**Figure S3. The ddG calculated from SPR versus** **log(Kd,mut/Kd,wt) from the DMS paper. The assays show contradict results on 4/9 of the mutants.**

**Table S2. List of the sample name on the SDS-PAGE figure corresponding to the mutants.**

| Sample name on the SDS-PAGE figure | Mutant |
| --- | --- |
| P101154-1 | RBD-Q498W |
| P101154-2 | RBD-Q498R |
| P101154-3 | RBD-T500W |
| P101154-4 | RBD-S477H |
| P101154-5 | RBD-Y505W |
| P101154-6 | RBD-T500R |
| P101154-7 | RBD-N501V |
| P101154-8 | RBD-Y489W |
| P101154-9 | RBD-Q493M |
| P101154-10 | RBD-L455A |

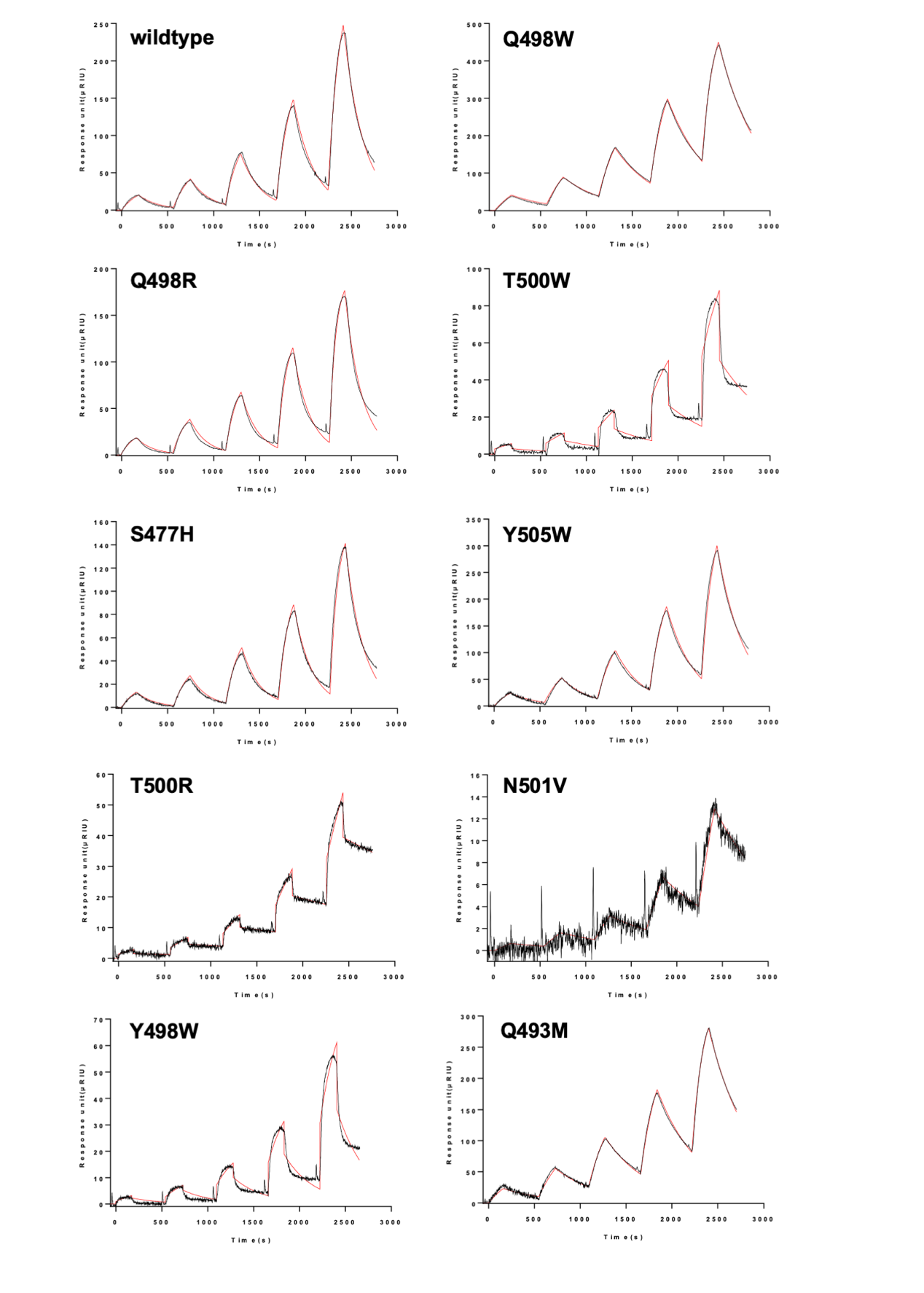

**Figure S4. Surface plasmon resonance sensorgram showing the binding kinetics for human ACE2 and immobilized SARS-CoV-2 S protein RBD wildtype / mutants. Data are shown as black lines, and the best fit of the data to a 1:1 binding model is shown in red.**

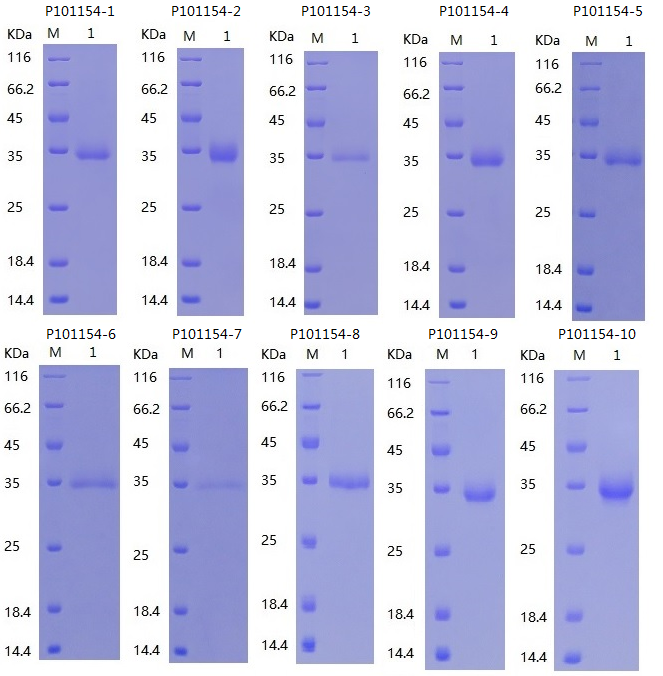

**Figure S5. SDS-PAGE result of the 10 spike protein RBD mutants. M: Protein marker. 1: Corresponding protein sample. The sample name and related mutant is shown in Table S2.**

**Table S3. SPR result of RBD-WT and 10 mutants.**

| Sample name | Protein concentration (mg/mL) | Ka (1/(M*s)) | Kd (1/s) | KD (M) |
| --- | --- | --- | --- | --- |
| RBD-WT | 0.25 | 3.6677E+04 | 9.048E-04 | 2.467E-8 |
| RBD-Q498W | 0.08 | 6.1757E+04 | 4.363E-04 | 7.065E-9 |
| RBD-Q498R | 0.35 | 9.3103E+04 | 1.081E-03 | 1.161E-8 |
| RBD-T500W | 0.06 | 1.4307E+04 | 3.117E-04 | 2.179E-8 |
| RBD-S477H | 0.19 | 7.3496E+04 | 1.021E-03 | 1.389E-8 |
| RBD-Y505W | 0.1 | 4.0716E+04 | 6.743E-04 | 1.656E-8 |
| RBD-T500R | 0.05 | 6.783E+03 | 8.253E-05 | 1.217E-8 |
| RBD-N501V | 0.05 | 1.623E+03 | 2.572E-04 | 1.585E-7 |
| RBD-Y489W | 0.1 | 1.5902E+04 | 6.189E-04 | 3.892E-8 |
| RBD-Q493M | 0.31 | 6.2167E+04 | 4.281E-04 | 6.886E-9 |
| RBD-L455A | 0.32 | 1.3389E+04 | 2.789E-04 | 2.083E-8 |

Ka: association (on) rate; Kd: dissociation (off) rate; KD: binding affinity constant
